# Bacterial mixed cultures followed mixed-order kinetics during mono-azo dye decolourization and degradation in batch studies

**DOI:** 10.1101/2023.04.08.536117

**Authors:** Kunal R. Jain, Digantkumar Chapla, Amita Shah, Datta Madamwar

**Author notes:** Corresponding Author (D. Madamwar), (K. Jain).

## Abstract

Over the years little has been focused on kinetic and thermodynamic aspect of bioremediation studies on dyes. In this study, detailed kinetic and thermodynamic analyses of results obtained during a primary study on a mono-azo dye: Reactive violet 5R (RV5), decolourization and degradation by bacterial mixed cultures SB4 were performed. Various kinetic models (zero-, first- and second-order behaviour and Monod model) were applied to understand the decolourization mechanism in batch process. Similarly, steady-state kinetics for determination of Michaelis-constant and maximal dye removal rate was derived. With increasing RV5 concentration, the specific decolourization rate had increased from 0.61mg/L/h to 1.38mg/L/h. The apparent kinetics of RV5 decolourization reaction followed mixed-order behaviour. The process was endothermic. The enthalpy and entropy values calculated to 45.6kJ/mol and 146J/mol/K, respectively, with an activation energy of 87kJ/mol. The Gibbs free energy change suggests that, decolourization reaction under experimental conditions was non-spontaneous below 20°C but was spontaneous at higher temperatures.

## 1.0 Introduction

For successful application of acclimatized bacterial mixed cultures (consortium) during the bioremediation process for dye containing industrial effluent, operation parameters (such as nutrient requirements, environmental factors, etc.) must be aptly recognized and studied. The primary aim of such studies is to enhance bioremediation potential of microorganisms or mixed cultures, before it can be considered for pilot scale or reactor studies or for on-field applications. Dafale et al. (2008), therefore have precisely noted that for developing an efficient bioremediation process for dye containing wastewaters, understanding related to kinetics of dye decolourization (*viz.* azoreduction, for reduction of azo bonds (–N=N–) of azo dye) and environmental factors affecting the dye degradation must be studied. Generally, studies related to decolourization of dye compounds were performed without further analysing and understanding the kinetic parameters affecting the decolourization and degradation process (Dafale et al., 2008; Lourenco et al., 2006).

It is normal practice to study the kinetics of a reaction step or a process of the entire metabolic pathway using a mathematical equation modelling method. The rationale behind these studies is to find an accurate model equation which can estimate the value of a rate of reaction that could be very close to the experimentally observed one. The selected kinetic model of a reaction system must consider all the essential parameters, is to be solved and the predicted behaviour should be compared with experimental derived results. If there is a lack of correlation between observed results and the behaviour predicted by the kinetic model, it usually implies the missing of few important parameters during the study. Further, the kinetic constants for each reaction/pathway are being estimated by fitting the model equation to the experimental results. The best fitted model thereafter, can be used for predicting the process performance under various operating conditions, reactor design, optimization, scale-up including control of the system (Houng and Liau, 2006; Bas et al., 2007).

Bioenergetics (thermodynamic) is an integral part of every biological reaction. Therefore, besides kinetic analysis, thermodynamic studies are also necessary to enhance the understanding of inherent energy changes and also facilitates in recognizing the energy requirement of the system during the bioremediation process. Since bioremediation is a biological process, energy (*viz.* temperature) requirement has binary function. Optimum temperature for microbial (i.e. cellular) growth often varies alongwith the requirement of substrate (dye decolourization and degradation) breakdown energy. Therefore, during the development of a bioremediation process, thermodynamic parameters must be equally considered.

It is obvious that the cellular as well as physiological behaviour of a bacterium would be significantly different in community than as a pure culture. So, it purely implies that kinetic behaviour alongwith thermodynamic constraints would vary considerably in community than in pure culture. Thus, considering the operational parameters during dye decolourization studies by the mixed cultures would not be adequate, their kinetic behaviour and thermodynamic performance must also be examined for predicting the successful completion of bioremediation processes. In our previous study (Jain et al., 2012), a bacterial mixed cultures SB4 were developed, its operational parameters (i.e. nutritional requirement and environmental (both biotic and abiotic factors)) for Reactive Violet 5R (RV5) decolourization and degradation was studied and optimized for getting maximum bioremediation potential by SB4.

Decolourization of RV5 by pure cultures or through bacterial mixed cultures is an enzymatic process, involving enzyme-substrate kinetics, which plays a vital role during decolourization (i.e. reduction of azo bond) and degradation of intermediates. It is known that, the reaction rate is a function of substrate concentration, hence, maximum substrate removal rate and half velocity constant would govern the dye decolourization and degradation process.

Therefore, kinetic analysis of RV5 decolourization was studied and various mathematical models were applied to understand the underlying decolourization mechanisms in batch process at flask level. Zero, first and second order kinetics with respect to the RV5 (i.e. substrate) concentration was applied along with Monod model which is most widely described in dye decolourization kinetic studies. Similarly steady-state kinetics for determination of Michaelis-Menten constant and maximal dye removal rate was also derived for the mixed cultures SB4. Moreover, decolourization of RV5 by bacterial mixed cultures requires a formation of enzyme-dye complex which must cross the energy barrier to catalyse the conversion of high molecular weight RV5 molecule into simpler compound. Thus, the study of bioenergetics (i.e. thermodynamics) of dye decolourization mechanism is an imperative to determine the spontaneity of the process and which was also studied using various established models.

## 2.0 Materials and Methods

### 2.1. Mixed cultures development and dye decolourization study

The development of mixed cultures SB4 (henceforth SB4), characterization for its nutrient requirement and effect of environmental parameters on Reactive violet 5R (RV5) dye decolourization (azoreduction) and its degradation was as described in our earlier study (Jain et al., 2012).

### 2.2 Kinetic study

#### 2.2.1. Monod kinetics

A kinetic model representing substrate removal, with simultaneous reduction and degradation of azo dyes can be traditionally expressed by deterministic kinetic model developed by Monod (*Equation1*). It has been frequently used for describing the rate of degradation of a substrate by living cells (Sponza and Isik., 2004),

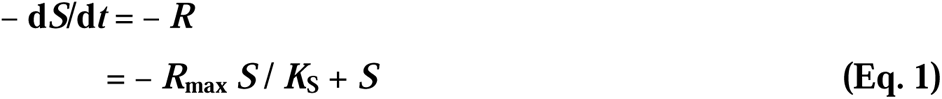

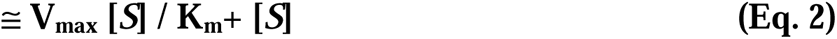

where *R*_max_ is maximum specific reduction rate (mg/L/h), *S* is substrate concentration, *K*_S_ is half saturation concentration (mg/L),

The *Equation 2* is generally known as Michaelis-Menten kinetics for monosubstrate reaction (where *K*_m_ ≅ *K*_S_, here *K*_S_ will be used in place of *K*_m_ which is also known as Michaelis-Menten constant). Upon linearizing *Equation 1*, the plot of *1/V* against *1/S*, produce straight line (i.e. Lineweaver-Burk plot) and would provide *V_max_*and *K*_m_.

#### 2.2.2. Determining the order of reaction for dye decolourization/degradation process

A general kinetic model for dye decolourization, including the effect of cell mass concentration, can be expressed by the following *Equation 3* (Lourenco et al., 2006),

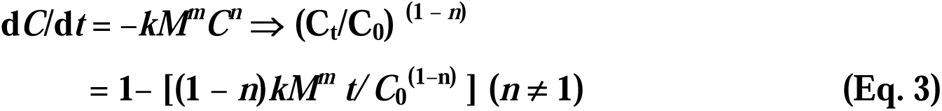

where *t* is the time (min), *M* and *m* are the cell mass concentration (mg dry cell mass/L) and its partial reaction order, respectively, *C_t_* and *n* are the dye concentration and its partial reaction order, respectively. The units of *k*, specific decolourization rate depends on the values of *m* and *n* [mg ^(1−^ *^n^*^)^ 1 ^(*m*^ ^+^ *^n^* **^−^** ^1)^/(mg dry cell mass*^m^* min)].

In a first-order reaction w.r.t. dye concentration, where *n* =1, with negligible cell growth or death (*M^m^* = constant), *Equation 3* can be simplified and rearranged to following

*Equation 4*:

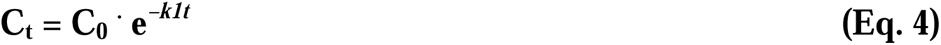

where *k_1_* is the first-order reduction rate (*1/h*) and its linear relationships are obtained by the plot of ln (*Ct*/*C_0_*) against time *t*, *C_t_* is concentration of dye at time *t* and *C_0_* is initial dye concentration.

To determine the order of reactions (i.e. zero, first or second), in order to estimate substrate degradation kinetics, experimental results obtained from batch studies from physico-chemcial parameters from earlier study (Jain et al., 2012) were plotted as: *C* verses time *t*, ln*C* verses time *t*, and *1/C* verses time *t* using *Equations 5, 6, 7* respectively.

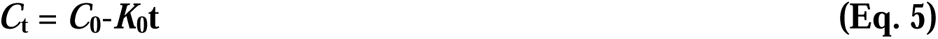

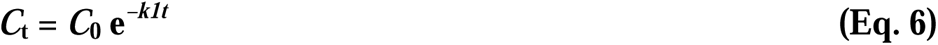

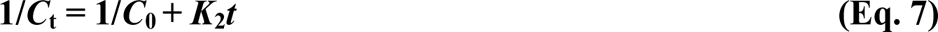

### 2.3 Thermodynamics (bioenergetics) of dye decolourization

#### 2.3.1 Activation energy

To correlate the temperature with the constant for the speed (velocity) of elementary reactions linearized Arrhenius equation was used/employed, which allows the determination of the activation energy alongwith the frequency factor for dye decolourization process, as expressed by *Equation 8* (Antelo et al., 2008),

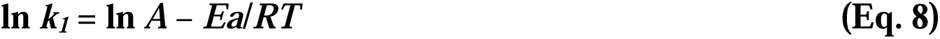

where *A* is the frequency factor, *Ea* is the activation energy, *R* is the gas constant (8.314 J/mol/K) and *T* is the absolute temperature (*K*). The values of constant *k_1_* measured at different temperatures were plotted against the reciprocal of the temperature and *Ea* was estimated from the slop of the plot (Annuar et al., 2009).

#### 2.3.2. Half-life estimation

The half-life *t_1/2_* (h) for first-order dye decolourization kinetics was estimated by *Equation 9* (Antelo et al., 2008),

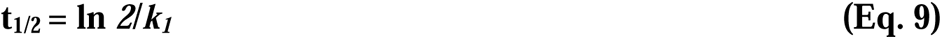

#### 2.3.3. Energetics parameters

The thermodynamic parameters such as enthalpy (*ΔH*) and entropy (*ΔS*) for RV5 decolourization were estimated using Van’t Hoff analysis (Annuar et al., 2009). It was assumed that dye decolourization process to be at an equilibrium state when the remaining RV5 concentration in the decolourization medium (solution) no longer changes with time. Therefore at equilibrium:

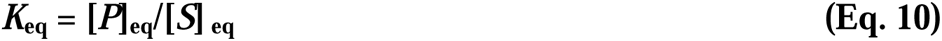

where *K*_eq_, is the apparent equilibrium constant, [*P*]_eq_ is the concentration of RV5 that has been decolourized at equilibrium and [*S*]_eq_ is RV5 concentration remaining at equilibrium.

The apparent equilibrium constant *K*_eq_ was measured at different temperatures. The temperature dependence of *K*_eq_ is expressed as Van’t Hoff equation as mentioned below:

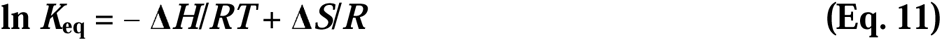

where *ΔH* is Van’t Hoff enthalpy (J/mol) and *ΔS* is the entropy (J/mol/K).

The above estimation assumes that decolourization of RV5 dye (i.e. reduction of azo bonds) occurs in an aqueous medium at a constant pressure without changes in the volume. Therefore, *ΔH* represents the heat transferred to a system at a constant pressure. At constant temperature and pressure, Gibbs free energy change (*ΔGm*, J/mol) for the reaction occurring at non-standard conditions was estimated using following equation:

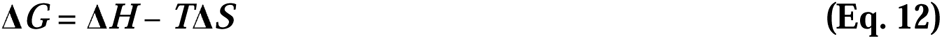

Standard free energy change (i.e. *ΔG*°, J/mol) was estimated using Gibbs–Helmholtz equation:

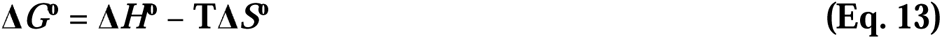

The *Equation 13* can be rearranged and also expressed as:

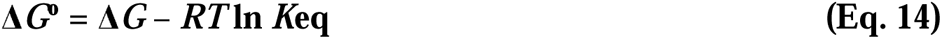

where *K_eq_* is the apparent equilibrium constant under standard conditions.

## 3.0 Results and Discussion

### 3.1 Determination of maximum substrate consumption rate and decolourization rate constant

Monod kinetics has been widely used for describing the rate of degradation of a substrate by living cells (Sponza and Isik, 2004). The dependence of specific decolourization rate (*V_max_*) on the dye (RV5) concentration was as observed from Figure 1. Initial specific dye decolourization rate was estimated for 100 to 1500 mg/L of RV5. At 400 mg/L, RV5 was decolourized at a specific decolourization rate of 0.61 mg/L/h, which increases to 1.38 mg/L/h for 1500 mg/L (Figure 1a). The low reduction rate can be acceptable, as it demonstrated in the previous study that it required nearly 24 h for complete reduction of 400 mg/L of RV5 by SB4 (Jain et al., 2012). The correlation between specific reduction rate and dye concentration can be studied by Michaelis–Menten kinetics. The kinetic constants, estimated from experimental results are 4.16 mg/L/h for *V_max_* and 2500 mg/L for *K_m_* (Figure 1b), for mixed cultures SB4. Such a high value of *K_m_*can be justified, as mixed culture SB4 has lower affinity for RV5, where it required nearly 18 h for complete decolourization of 100 mg/L dye (Jain et al., 2012). In the previous study on decolourization of textile waste water by pure culture *Staphylococcus cohnii* by Yan et al., (2012), found *V_max_* as 109.8 mg/g cell/h and *K_m_* as 585.7 mg/L. Similarly, Chang et al., (2001), estimated the kinetic constants for azo dye degradation by *Pseudomonas luteola* as 12.0 mg/g cell/h for *V_max_* and 156 mg/L for *K_m_*.

**Figure 1:**
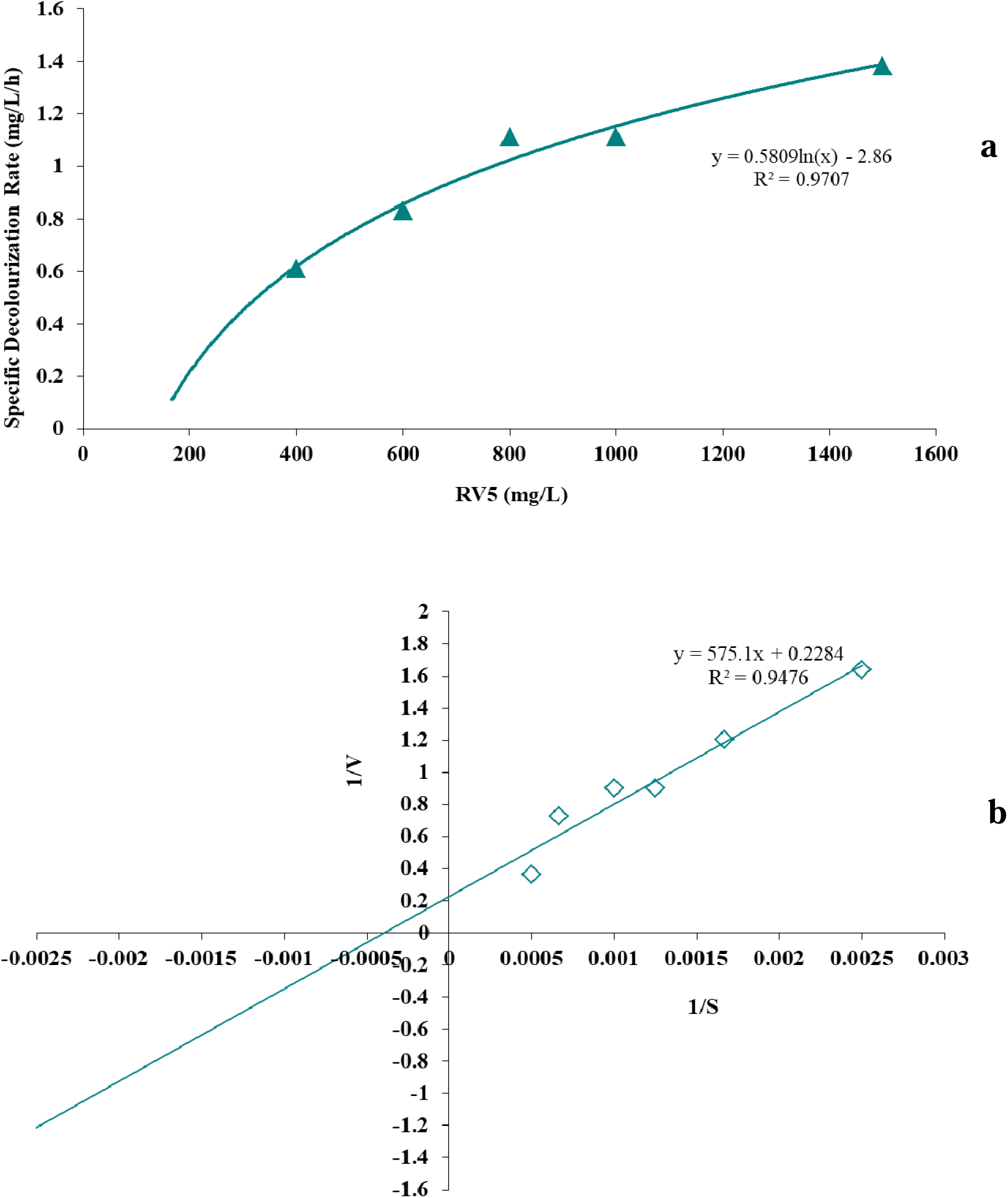
Relation between RV5 concentration and specific decolourization rate**. (a)** The effect of substrate (dye) concentration on the specific decolourization rate of RV5 by mixed culture SB4. **(b)** Lineweaver-Burk plot for estimating kinetic constants.

### 3.2 Determining the order of the reaction

Results obtained from batch studies for substrate utilization (i.e. RV5 decolourization) were plotted as mentioned above. Kinetic coefficients and correlation coefficients relevant to zero, first and second orders (*k*_0_, *k*_1_ and *k*_2_ values) were estimated through straight lines. The experimental results obtained during this study revealed surprising observations. Substrate (RV5 decolourization) removal reaction did not follow any particular order and it was evident that dye decolourization by SB4followed mixed order kinetics and depended on the concentration of RV5.

#### 3.2.1 Dye (substrate) concentration

At lower RV5 concentrations (100/200 mg/L), for SB4, the regression coefficient was 1.0 for all three reaction orders as evident from Table 1 and Figure 2. At higher concentrations (400 to 1500 mg/L) dye decolourization reaction found to follow zero order kinetics (since linear relationship was comparatively higher, R^2^ >0.99, between RV5 concentration *C* and time *t*). The study further revealed that, reaction constant for zero order kinetics gradually increased in the range of 21.7 to 60 1/h. The earlier studies with respect to substrate utilization have found the prevailing of zero order kinetics (Dubin and Wright, 1975; Brown, 1981) as well as first order kinetics (Xie et al., 2020; Carliell et al., 1995; van der Zee et al., 2001; Wuhrmann et al., 1980; Weber et al., 1987; Weber, 1991).

**Figure 2:**
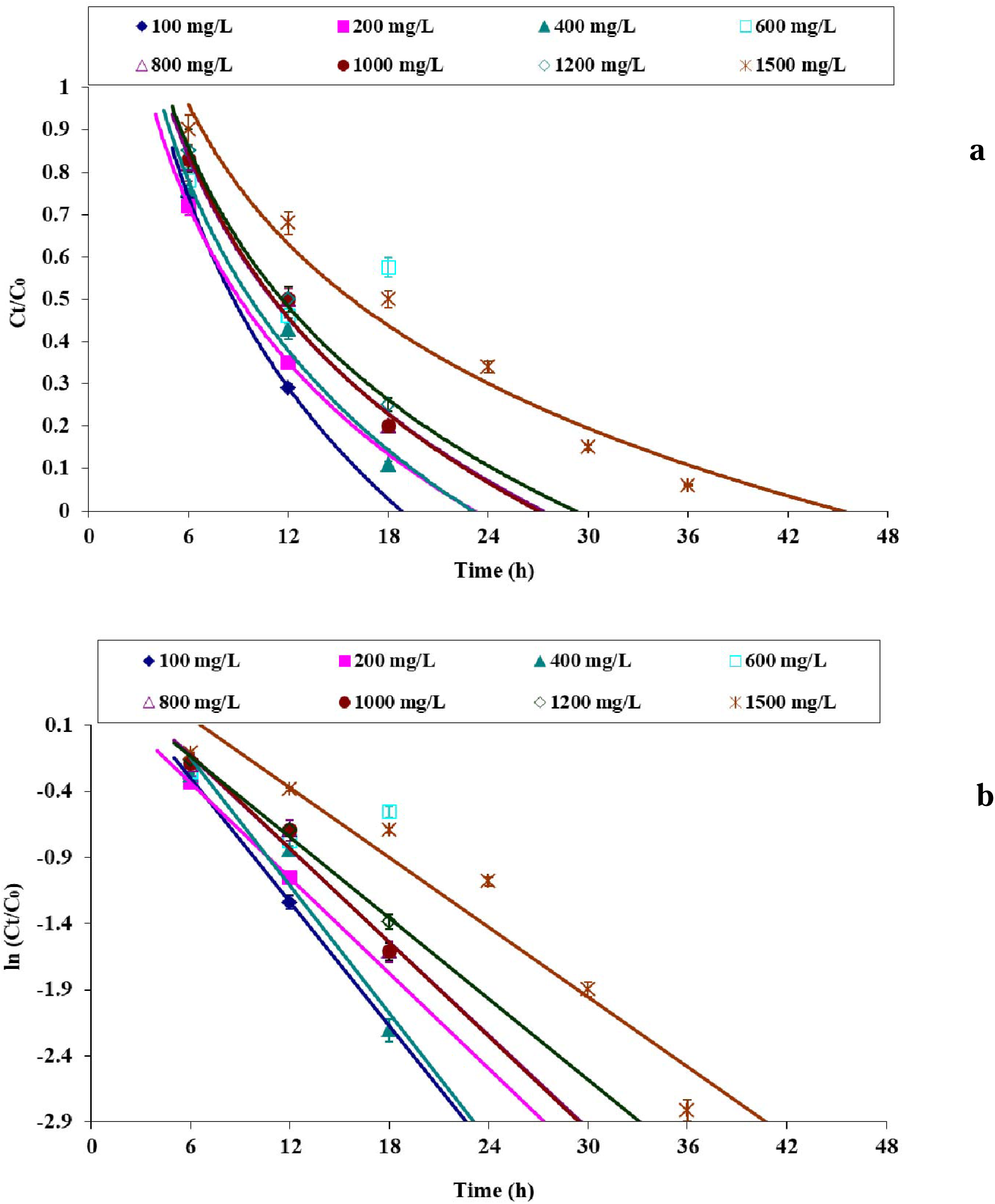
Decay curves for various RV5 concentration; **(a)** Plot of *C_t_/C_0_* against time for decolourization of monoazo RV5 dye by mixed cultures SB4, showing decrease in dye concentration with reaction following zero order kinetics at higher concentration (≥ 400 mg/L). **(b)** Linearized plot of ln(*C_t_/C_0_*) against time showing decreasing in dye concentration, where reaction followed zero order kinetics at higher concentration (≥ 400 mg/L).

**Table 1:** Kinetic constants for substrate utilization (i.e. RV5 decolourization) by mixed cultures SB4 under experimental conditions in minimal medium

| Kinetics | Constant | 100<br>mg/L | 200<br>mg/L | 400<br>mg/L | 600<br>mg/L | 800<br>mg/L | 1000<br>mg/L | 1200<br>mg/L | 1500<br>mg/L |
| --- | --- | --- | --- | --- | --- | --- | --- | --- | --- |
| Zero<br>Order | $k_0$<br>(mg/L/h) | 7.5000 | 12.3000 | 21.7000 | 31.5000 | 41.3000 | 52.5000 | 60 | 51 |
| | $R^2$ | 1 | 1 | 0.9990 | 0.9990 | 0.9990 | 0.9990 | 0.9900 | 0.9950 |
| First<br>Order | $k_1$<br>(1/h) | 0.1561 | 0.1202 | 0.1611 | 0.1374 | 0.1176 | 0.1186 | 0.1020 | 0.1028 |
| | $R^2$ | 1 | 1 | 0.9468 | 0.9587 | 0.9711 | 0.9731 | 0.9942 | 0.8319 |
| Second<br>Order | $k_2$<br>(L/mg/h) | 0.0035 | 0.0012 | 0.0016 | 0.0007 | 0.0004 | 0.0003 | 0.0002 | 0.0004 |
| | $R^2$ | 1 | 1 | 0.8456 | 0.8703 | 0.8969 | 0.8989 | 0.9453 | 0.6095 |

SB4 does not found to follow either first or second order kinetics with respect to substrate (RV5) concentration. However the analysis on dye decolourization constant for higher RV5 concentration (400 to 1200 mg/L), found to decrease the reaction constant from 0.1611 to 0.1020 mg/L/h for first order kinetic. And for second order kinetics it decreased gradually from 0.0016 to 0.0002 mg/L/h with increasing RV5 concentrations from 200 mg/L to 1200 mg/L (Table 1). Similar decrease in rate constant with increasing substrate (dye) concentration was found in few of the studies (Cao et al., 2019; Eskandari et al., 2019).

#### 3.2.2. Carbon source

Similarly, for utilization of various carbon sources, different order kinetics was also applied. The analysis indicated that SB4 followed zero order kinetics for sucrose, starch, lactose and pyruvate and that for glucose it was mixed order kinetics (Table 2). The rate constant (*k*_0_) gradually increased for lactose, pyruvate, starch and sucrose from 1.9464 to 2.3274 mg/L/h respectively, while for glucose, considering the zero order kinetics, it was maximum at 6.8333 mg/L/h. These kinetic results have been corroborated with the results obtained in the primary study on RV5 decolourization (Jain et al., 2012). Since glucose was the preferable carbon source during RV5 decolourization by SB4 the rate constant has to highest amongst the carbon source used in the study. The analysis also indicated that RV5 decolourization efficiency increased from lactose to sucrose (section 2.2.--) which can evidently observed with increase in rate constant *k*_0_.

**Table 2:** Kinetic constants for various carbon sources used during decolourization study by mixed cultures SB4 under experimental conditions in minimal medium

| Kinetics | Constant | Glucose | Lactose | Sucrose | Pyruvate | Starch |
| --- | --- | --- | --- | --- | --- | --- |
| Zero | $k_0$ | 6.8333 | 1.9464 | 2.3274 | 2.0159 | 2.2877 |
| Order | (mg/L/h) |  |  |  |  |  |
| | $R^2$ | 0.9968 | 0.9539 | 0.8706 | 0.9717 | 0.8052 |
| First | $k_1$ | 0.1000 | 0.0374 | 0.0803 | 0.0478 | 0.0680 |
| Order | (1/h) |  |  |  |  |  |
| | $R^2$ | 1 | 0.9196 | 0.8385 | 0.9634 | 0.7541 |
| Second | $k_2$ | 0.0017 | 0.0008 | 0.0033 | 0.0014 | 0.0032 |
| Order | (L/mg/h) |  |  |  |  |  |
| | $R^2$ | 1 | 0.8477 | 0.6760 | 0.8717 | 0.5751 |

For glucose concentration the reaction kinetic model varies with varying glucose concentration (Table S1). At higher glucose concentration (≥ 2 g/L) SB4 followed first order kinetic model.

#### 3.2.3. Alternate electron acceptors

Decolourization of RV5 is an oxido-reduction process. Along with dissolved oxygen, nirates and carbonates act as alternate electron acceptor during dye decolourization. In the presence of molecular oxygen, dye decolourization reaction followed second and zero order (Table 3). RV5 decolourization without any external electron acceptor followed nearly mixed order reaction kinetics. Nitrates followed first and second order reaction kinetics, while carbonates followed second order kinetics. The reaction constant for different salts of nitrates (potassium, sodium and ammonium salts) gradually decrease, while the rate constant *k_2_,* for carbonates was estimated to be 0.0001 L/mg/h.

**Table 3:**
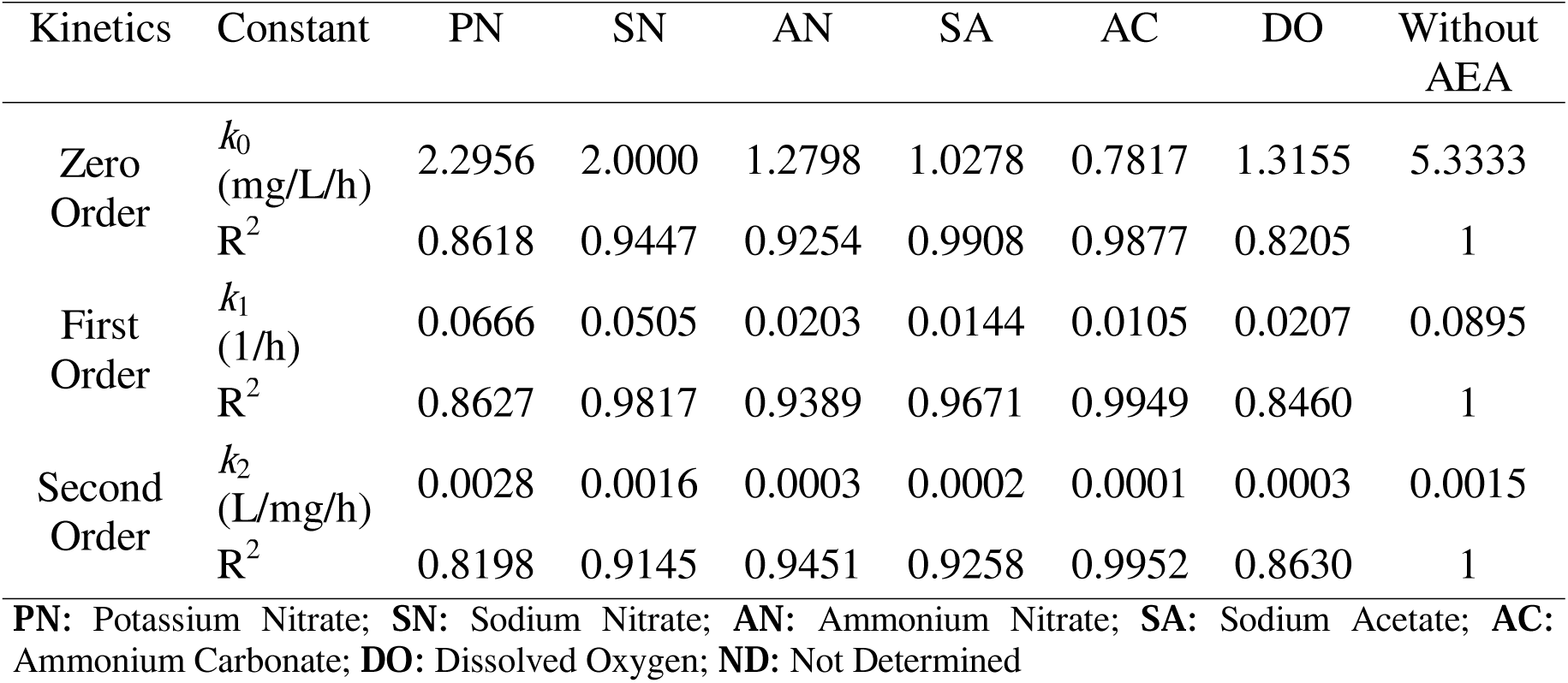
Kinetic constants for various source of electron acceptors used during decolourization study by mixed cultures SB4 under experimental conditions in minimal medium

#### 3.2.4. Nitrogen source

Similarly, w.r.t. nitrogen source, SB4 followed zero order reaction kinetics for urea and peptone, and mixed order reaction kinetics for yeast extract (Table 4). For tryptone it was first order reaction kinetics. Correspondingly, for yeast extract concentration, the estimation suggests that for ≥ 1g/L yeast extract concentration, it was mixed order reaction kinetics (Table S2).

**Table 4:** Kinetic constants for various nitrogen sources used during decolourization study by mixed cultures SB4 under experimental conditions in minimal medium

| Kinetics | Constant | Yeast<br>Extract | Peptone | Urea |
| --- | --- | --- | --- | --- |
| Zero<br>Order | $k_0$<br>(mg/L/h) | 1.5655 | 1.2639 | 2.1806 |
| | $R^2$ | 0.5598 | 0.5722 | 0.9083 |
| First<br>Order | $k_1$<br>(1/h) | 0.1076 | 0.0292 | 0.0522 |
| | $R^2$ | 1 | 0.5266 | 0.8989 |
| Second<br>Order | $k_2$<br>(L/mg/h) | 0.0018 | 0.0008 | 0.0016 |
| | $R^2$ | 1 | 0.4011 | 0.8486 |

**Table 5:**
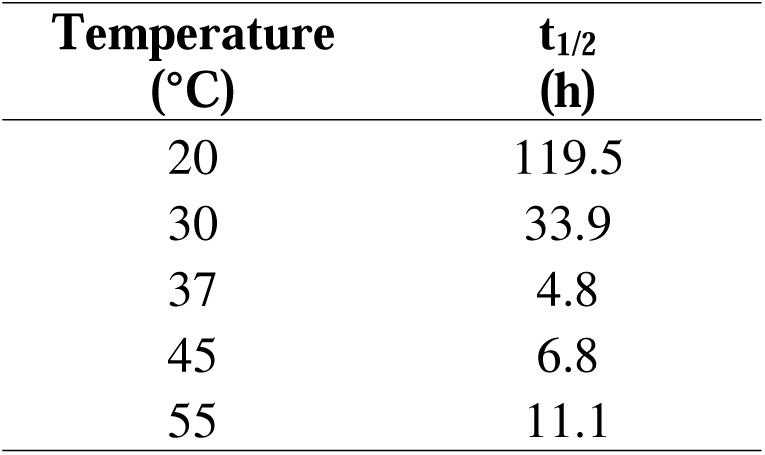
Half-life values for RV5 decolourization by mixed cultures SB4 at different temperature under experimental conditions in minimal medium

#### 3.2.5. pH and Temperature

The results from Table S3 revealed that RV5 decolourization by SB4, followed first order reaction kinetics between pH 5.0 and 6.5. At pH 7.0, it was a mixed order reaction which was an optimum pH for RV5 decolourization. For higher pH 8.0/9.0 it followed first and zero order reaction kinetics. The rate constants increased gradually while reaching to maximum value at optimum pH (i.e. 7.0) and then decreased at higher pH. Decolourization of RV5 by mixed cultures SB4 followed mixed order kinetics, at 37°C, while for higher temperature 45/55°C, it followed first order reaction kinetics (Table S4). With the increase in temperature (from 20 and 37°C), as expected first order kinetic constant, *k_1_* also increased, and at higher temperature (40 °C), obviously kinetic constant decreased. Dafale et al., (2008), with their bacterial consortium, capable of decolourizing Remazol black-B dye, observed that consortium followed first order reaction kinetics between temperatures 20 and 37°C.

#### 3.2.6. Salt (NaCl) concentration

The results from the Table S5 revealed that decolourization of RV5 by SB4 followed mixed order kinetics medium devoid of salts (NaCl) followed mixed order reaction kinetics, while azoreduction by consortia AIE2 and AVIE2 in absence of salt followed first and zero order reaction kinetics respectively and having maximum value of rate constants. Mixed cultures SB4 at salt concentration of 2 g/L, followed mixed order kinetics, at 4 g/L and 10 g/L it was zero and first order and at higher salt concentration reaction followed first order kinetics.

### 3.3. Thermodynamics of dye decolourization

#### 3.3.1. Activation energy and half-life

During decolourization study temperature was found to have profound effect on microaerophilic reduction of azo bonds of RV5 (Jain et al., 2012). The efficiency of RV5 decolourization increases with increase in temperature from 20 to 42°C and thereafter declined, which depends on activation energy of the reaction (Jain et al., 2012; Dafale et al., 2008). The obtained results with respect to the temperature, evidently suggest that RV5 decolourization by the mixed cultures SB4 followed first order kinetics. Therefore, the apparent activation energy (*Ea*) for decolourization of RV5 (i.e. reduction of azo bonds of RV5) was estimated from Arrhenius plots as shown in Figure 3, and was estimated to be 87 kJ/mol.

**Figure 3:**
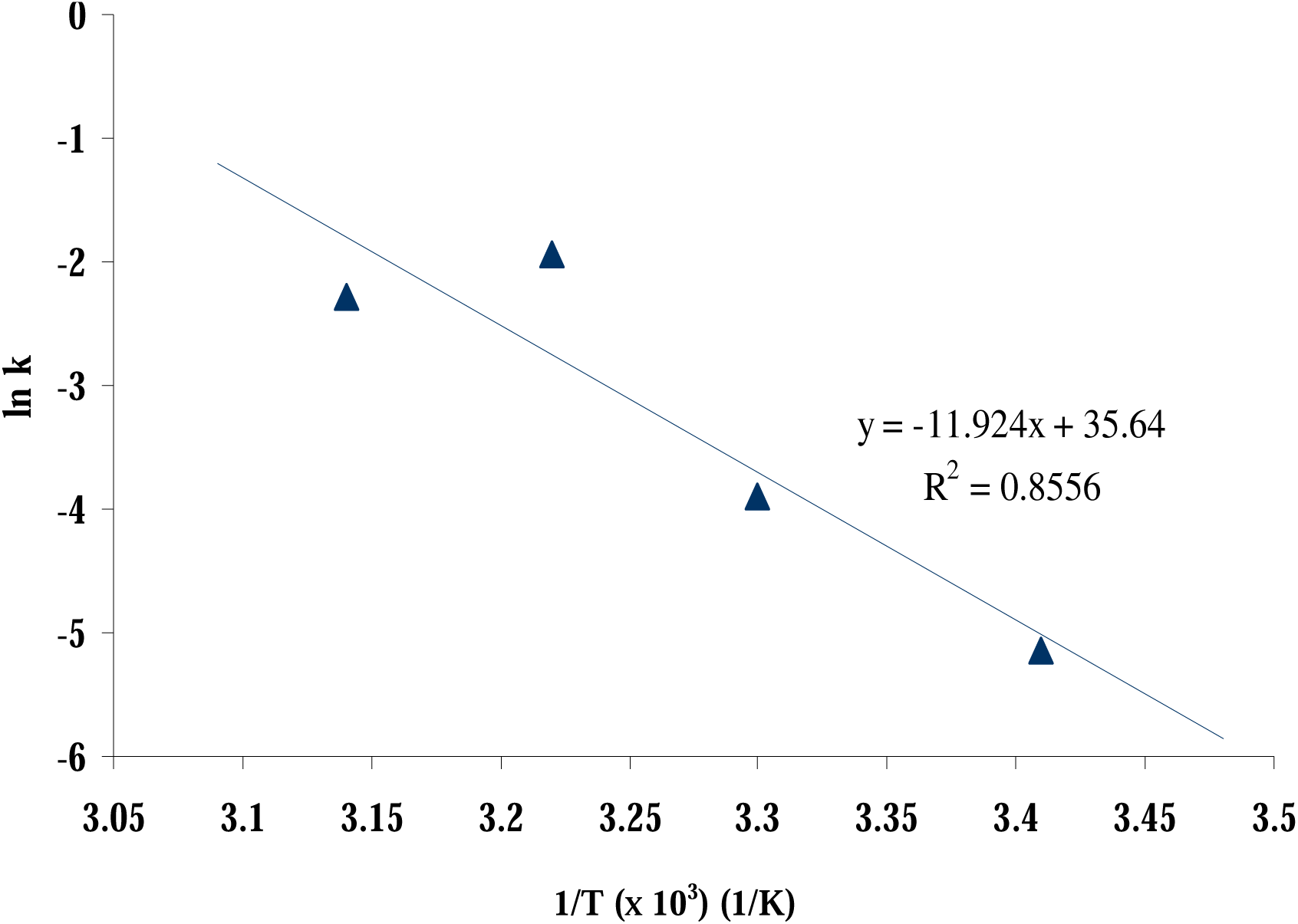
Arrhenius plot of rate constant against temperature for decolourization of RV5 by mixed cultures SB4 under experimental conditions.

Since the plot produced high degree of linearity along with regression coefficient of > 0.95, the obtained value offered reliable estimation of activation energy. Moreover, through different studies the activation energy values for enzyme-catalyzed reactions was estimated to be in the range of 16 to 84 kJ/mol (Annuar et al., 2009). Therefore, the *Ea* for SB4-RV5 reaction can be reasonably accepted. During their study, Dafale et al., (2008) estimated the activation energy of 11.67 kcal/mol (i.e. 48.82 kJ/mol) for decolourization of Remazol black-B by a bacterial consortium in a two-stage continues reactor. In another study, degradation of Trypan blue by bacterium *Pycnopotus sanguineus*, required the activation energy equivalent to 23 kJ/mol (Annuar et al., 2009). Alongwith the activation energy, half-life for reduction of azo bonds (i.e. dye decolourization) of RV5 was estimated, which was 4.8 h, under given decolourization conditions for SB4 (Table 6). The calculated half-life was corroborated with the primary results observed for RV5 decolourization i.e. it required 18 h for complete decolourization/degradation of RV5 (200 mg/L) by SB4 (Jain et al., 2012). Higher value of half-life for SB4-RV5 reaction therefore was agreeable. Results from the Table 6, revealed that half-life values gradually decreases and is minimal at optimal temperature of 37°C. At higher temperatures (< 42°C) half-life values again increases as the rate of decolourization decreases.

**Table 6:**
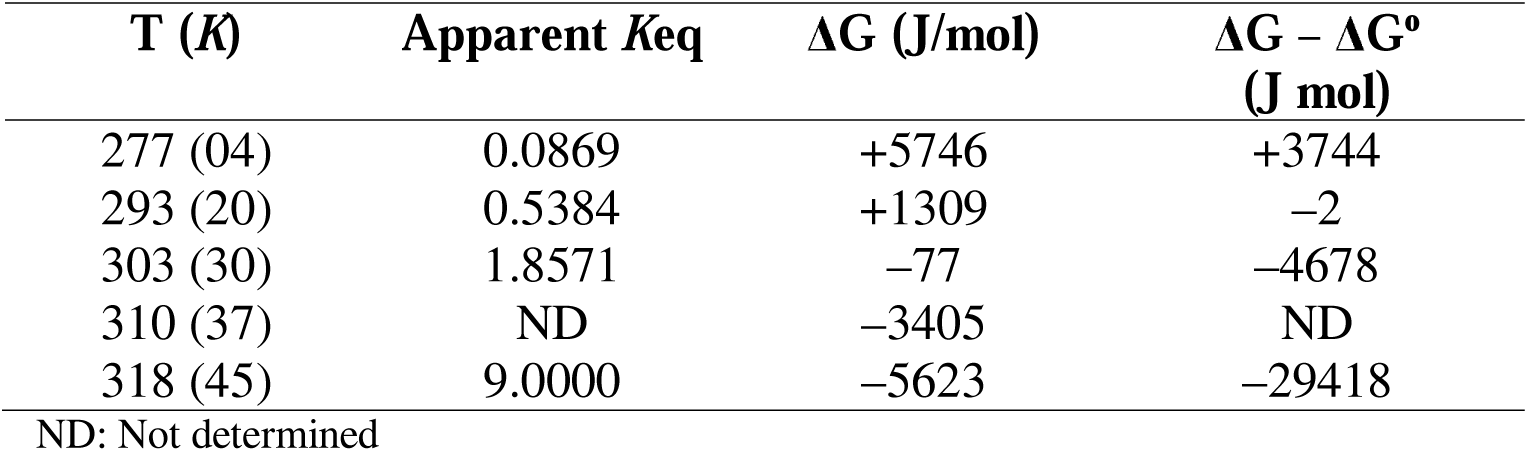
Gibbs free energy and Gibbs free energy change for the reaction catalysing the reduction of azo bonds (i.e. decolourization of RV5) by mixed cultures SB4 at different temperatures

#### 3.3.2. Spontaneity of the process

Thermodynamic parameters (i.e. enthalpy and entropy) essentially provide the inherent energy changes associated with decolourization of RV5 (reduction in azo bonds) by bacterial mixed cultures (Ahmad et al., 2014) To determine the spontaneity of the process and system, the apparent enthalpy (Δ*H*) as well as entropy (Δ*S*) for RV5 decolourization was estimated using Van’t Hoff plot (Figure 4). In the analysis for the determination of above thermodynamic parameters, the implicit assumption, i.e. a closed system at a constant pressure and volume were applied for dye decolourization reaction.

**Figure 4:**
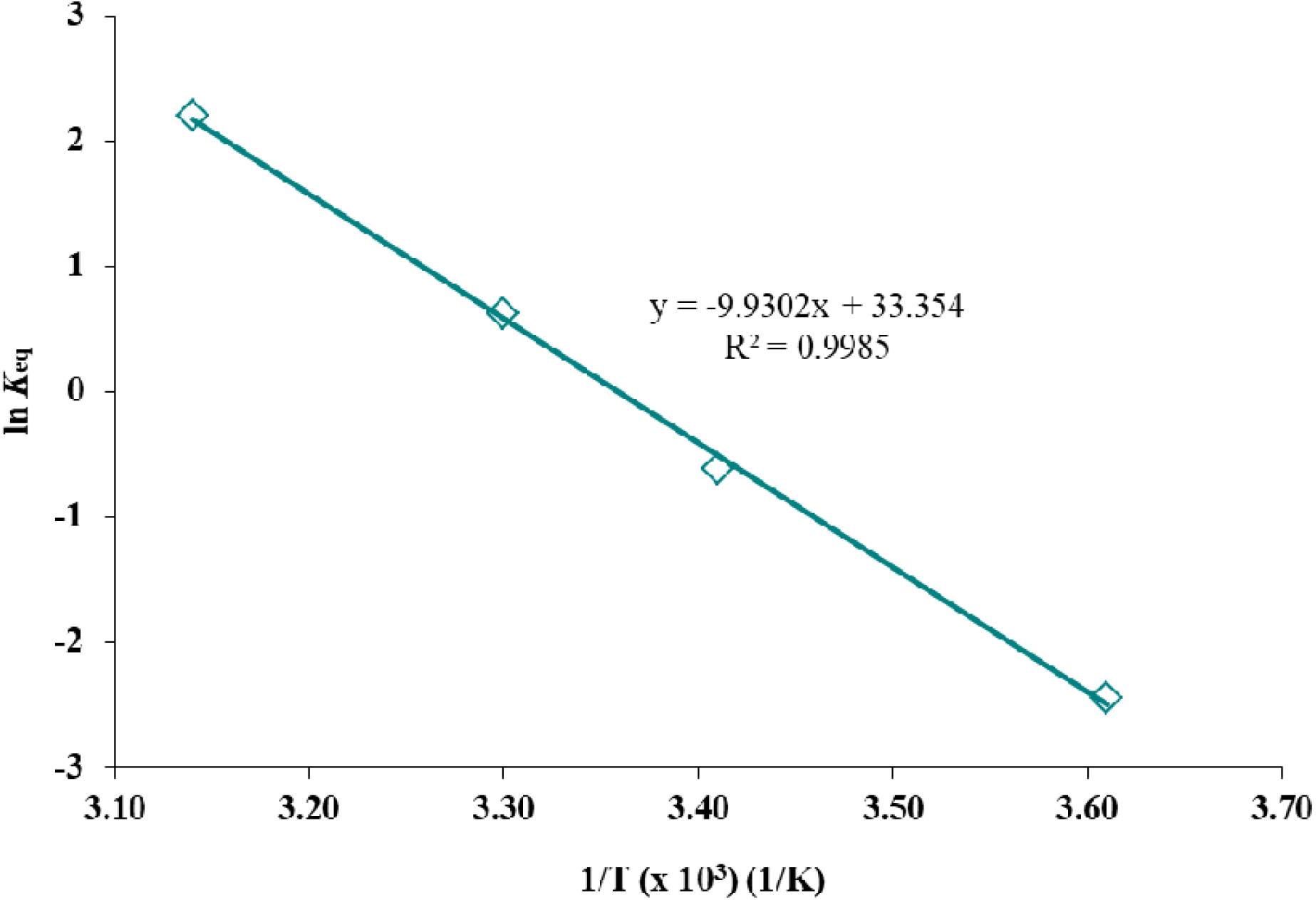
Van’t Hoff plot of equilibrium constant against temperature for decolourization of RV5 by mixed cultures SB4 under experimental conditions.

The enthalpy (*ΔH*) for decolourization (i.e. reduction of azo bonds of RV5) by SB4 was estimated to be 82.55 kJ/mol and entropy (Δ*S*) was 277.30 J/mol K. The positive values of enthalpy and entropy suggest an endothermic reduction of azo bonds, consequently was also accompanied by an increased disorder in the system as the molecular structure of RV5 decomposed (Annuar et al, 2009). The primary study also corroborates with the positive values of estimated enthalpy, which explains the increase in rate of RV5 decolourization with increase in temperature (Jain et al., 2012).

Besides above parameters, an important element of free energy change i.e. Δ*G*: Gibbs free energy, for decolourization of RV5 along with its degradation at various temperatures was also estimated using Van’t Hoff plot (Table 6). The results evidently suggested that the decolourization (and also further degradation) of RV5 by SB4 under studied conditions was non-spontaneous from temperature 20°C and below, as indicated by positive value of Δ*G*. But with the increasing in temperature form 30°C and above, the negative values of Δ*G* indicated the spontaneity of the reaction. The array of results obtained assumes that mass transfer of dye compound was not a limiting factor for its decolourization and degradation (Annuar et al., 2009).

The spontaneity of the decolourization of RV5 under experimental conditions was further validated by plotting Gibbs free energy (Δ*G*) for dye decolourization reaction at different temperatures. The results from the Figure 5, revealed that at 25°C, Δ*G* becomes zero and below 25°C, Δ*G* was positive, indicating the non-spontaneity of dye decolourization. However above 25°C, Δ*G* becomes negative, indicating that decolourization of RV5 by SB4 was favoured at an above 25°C. These results also similar with the observed values of Δ*G* estimated through *Equation 12*, (and as shown in Table 6), where Δ*G* was negative at 30°C.

**Figure 5:**
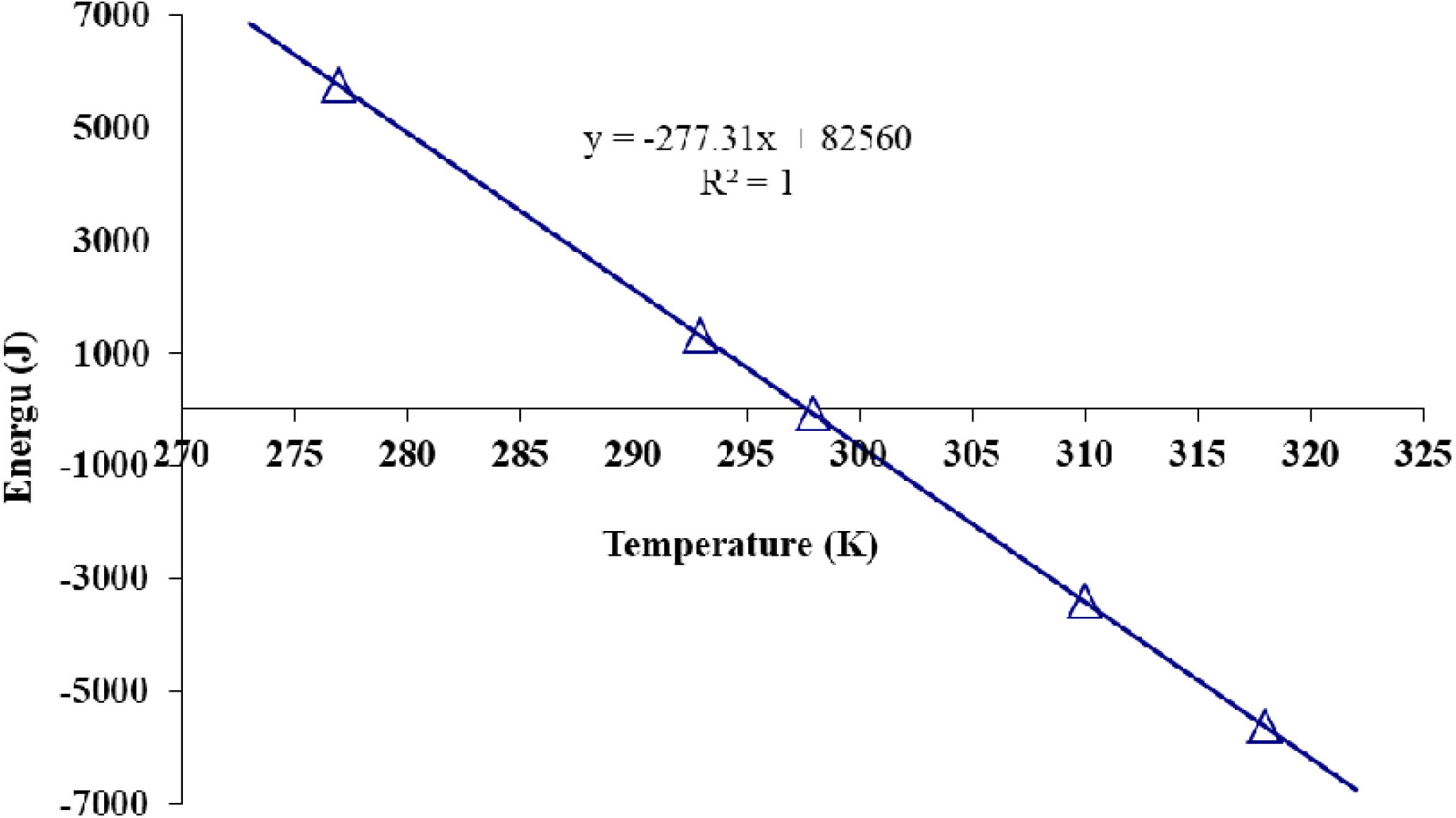
Gibbs free energy change for the reaction catalysing the decolourization of RV5 dye by mixed cultures SB4, under experimental condition as a function of temperature.

### 3.4 Theoretical implications

Decolourization of Reactive violet 5R using bacterial mixed cultures SB4 is a biological and enzymatic process, actively involving enzyme-substrate kinetics that play a pivotal role in dye degradation. The rate of reaction of ES-complex is a function of substrate concentration, maximum substrate removal rate, half velocity constants that governs the dye decolourization and degradation process. Though decolourization of mono-azo dye have been reported to follow first order kinetics, few of them demonstrated to follow zero-order or even half-order kinetics. As noted above, RV5 decolourization by SB4 at lower dye concentration, the reaction rate was linear which turned to mixed ordered at higher concentration and was which was a very unique observation in dye decolourization studies. And with increase in dye concentration the kinetic constant also increased with increasing in decolourization rate, which normally does not in dye decolourization studies.

During primary study (Jain el at., 2012), the dependence of RV5 decolourization on co-substrates (carbon and nitrogen sources) was evident. The mixed-order kinetics (i.e. SB4 must have followed first-order kinetics at some phase of RV5 decolourization) estimated in this study also supports the dependence on co-substrates. With the positive value of standard enthalpy (Δ*H°*) change, RV5 decolourization (and further degradation) was endothermic. While the standard entropy (Δ*S°*) change also estimated as positive, the disorder in the system (i.e. increasing the randomness during the degradation of RV5 under batch mode) might have also increased. The analysis was evidently corroborated with primary study on RV5, where four RV5 degraded intermediates were detected, i.e. molecular structure of dye compound was metabolized (degraded) into smaller compounds increasing the disorder of the system.

## 4.0 Conclusion

This ancillary study is one of the very few studies in which various kinetic models along with thermodynamics were applied, to understand the behaviour between dye decolourization process and bacterial cells in the mixed form. The SB4 followed mixed order kinetics primarily at higher concentration of dye. The applied thermodynamics suggested that, decolourization of RV5 using mixed cultures was endothermic and became spontaneous above 25°C. The observed results therefore is significant, which has provided an essential understanding about behaviour of bacteria in community, that can be used to develop an efficient bioremediation strategy for mixed culture bioremediation of dye compounds.

## Acknowledgement

Authors are grateful Department of Biotechnology (DBT), Ministry of Science and Technology, New Delhi, India for financial support. KJ duly acknowledges Council of Scientific and Industrial Research (CSIR), New Delhi, India for financial support.

## Supplementary Data

**Table S1:** Kinetic constants varying glucose concentration utilize during decolourization of RV5 by mixed cultures SB4 under experimental conditions in minimal medium

| Kinetics | Constant | 0.1 g/L | 0.5 g/L | 1 g/L | 2 g/L | 3 g/L | 5 g/L |
| --- | --- | --- | --- | --- | --- | --- | --- |
| Zero | $k_0$ | 0.3075 | 1.8214 | 6.6667 | 2.0476 | 1.7262 | 2.0317 |
| Order | (mg/L/h) |  |  |  |  |  |  |
| | $R^2$ | 0.8727 | 0.8807 | 1 | 0.7915 | 0.7334 | 0.7951 |
| First | $k_1$ | 0.0039 | 0.0438 | 0.1115 | 0.0992 | 0.0598 | 0.0801 |
| Order | (1/h) |  |  |  |  |  |  |
| | $R^2$ | 0.8751 | 0.8694 | 1 | 0.8902 | 0.7910 | 0.9590 |
| Second | $k_2$ | 5E-05 | 0.0013 | 0.0019 | 0.0150 | 0.0031 | 0.0064 |
| Order | (L/mg/h) |  |  |  |  |  |  |
| | $R^2$ | 0.8755 | 0.7266 | 1 | 0.4353 | 0.7033 | 0.7889 |

**Table S2:** Kinetic constants varying yeast extract concentration utilize during decolourization of RV5 by mixed cultures SB4 under experimental conditions in minimal medium

| Kinetics | Constant | 0.1 g/L | 0.5 g/L | 1 g/L | 2 g/L | 3 g/L | 5 g/L |
| --- | --- | --- | --- | --- | --- | --- | --- |
| Zero Order | $k_0$<br>(mg/L/h) | 2.5377 | 2.3552 | 6.6667 | 6.6667 | 6.4167 | 6.4167 |
| | $R^2$ | 0.9036 | 0.8956 | 0.9992 | 0.9796 | 0.9487 | 0.8176 |
| First Order | $k_1$<br>(1/h) | 0.0920 | 0.0998 | 0.1074 | 0.1635 | 0.2014 | 0.3996 |
| | $R^2$ | 0.8296 | 0.8014 | 1 | 1 | 1 | 1 |
| Second Order | $k_2$<br>(L/mg/h) | 0.0080 | 0.0133 | 0.0019 | 0.0035 | 0.0051 | 0.0216 |
| | $R^2$ | 0.4801 | 0.3405 | 1 | 1 | 1 | 1 |

**Table S3:**
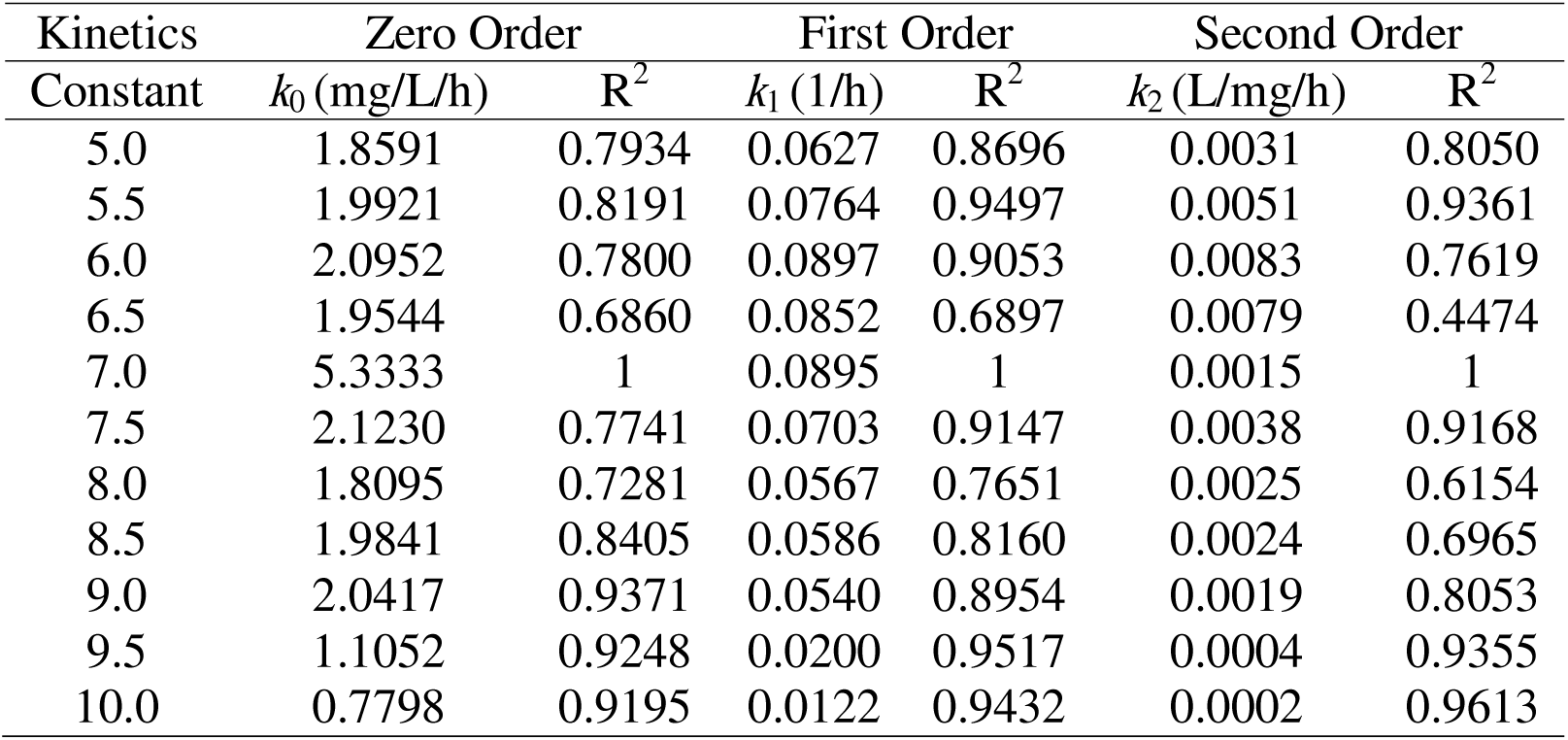
Kinetic constants for decolourization of RV5 by mixed cultures SB4 at different pH, under experimental conditions in minimal medium

| Kinetics | Zero Order |  | First Order |  | Second Order |  |
| --- | --- | --- | --- | --- | --- | --- |
| Constant | $k_0$ (mg/L/h) | $R^2$ | $k_1$ (1/h) | $R^2$ | $k_2$ (L/mg/h) | $R^2$ |
| 5.0 | 1.8591 | 0.7934 | 0.0627 | 0.8696 | 0.0031 | 0.8050 |
| 5.5 | 1.9921 | 0.8191 | 0.0764 | 0.9497 | 0.0051 | 0.9361 |
| 6.0 | 2.0952 | 0.7800 | 0.0897 | 0.9053 | 0.0083 | 0.7619 |
| 6.5 | 1.9544 | 0.6860 | 0.0852 | 0.6897 | 0.0079 | 0.4474 |
| 7.0 | 5.3333 | 1 | 0.0895 | 1 | 0.0015 | 1 |
| 7.5 | 2.1230 | 0.7741 | 0.0703 | 0.9147 | 0.0038 | 0.9168 |
| 8.0 | 1.8095 | 0.7281 | 0.0567 | 0.7651 | 0.0025 | 0.6154 |
| 8.5 | 1.9841 | 0.8405 | 0.0586 | 0.8160 | 0.0024 | 0.6965 |
| 9.0 | 2.0417 | 0.9371 | 0.0540 | 0.8954 | 0.0019 | 0.8053 |
| 9.5 | 1.1052 | 0.9248 | 0.0200 | 0.9517 | 0.0004 | 0.9355 |
| 10.0 | 0.7798 | 0.9195 | 0.0122 | 0.9432 | 0.0002 | 0.9613 |

**Table S4:** Kinetic constants for decolourization of RV5 by mixed cultures SB4 at different temperature, under experimental conditions in minimal medium

| Kinetics | Zero Order |  | First Order |  | Second Order |  |
| --- | --- | --- | --- | --- | --- | --- |
| Constant | $k_0$ (mg/L/h) | $R^2$ | $k_1$<br>(1/h) | $R^2$ | $k_2$<br>(L/mg/h) | $R^2$ |
| 20°C | 0.8214 | 0.9735 | 0.0109 | 0.9768 | 0.0001 | 0.9760 |
| 30°C | 1.9921 | 0.9773 | 0.0442 | 0.9414 | 0.0012 | 0.8159 |
| 37°C | 5.3333 | 1 | 0.0895 | 1 | 0.0015 | 1 |
| 45°C | 2.0337 | 0.8704 | 0.0680 | 0.9613 | 0.0036 | 0.9076 |
| 55°C | 1.6647 | 0.9473 | 0.0337 | 0.9784 | 0.0008 | 0.9367 |

**Table S5:** Kinetic constants for decolourization of RV5 by mixed cultures SB4 at different salt (NaCl) concentration, under experimental conditions in minimal medium

| Kinetics | Zero Order |  | First Order |  | Second Order |  |
| --- | --- | --- | --- | --- | --- | --- |
| Constant | $k_0$ (mg/L/h) | $R^2$ | $k_1$ (1/h) | $R^2$ | $k_2$<br>(L/mg/h) | $R^2$ |
| Without NaCl | 6.3333 | 1 | 0.1225 | 1 | 0.0025 | 1 |
| 1 g/L | 5.6667 | 1 | 0.1045 | 1 | 0.002 | 1 |
| 2 g/L | 5.6667 | 1 | 0.1007 | 1 | 0.0018 | 1 |
| 3 g/L | 5.0000 | 0.9908 | 0.1222 | 0.9884 | 0.0036 | 0.9197 |
| 4 g/L | 4.8333 | 0.9937 | 0.1222 | 0.9884 | 0.0036 | 0.9197 |
| 5 g/L | 3.5000 | 0.9866 | 0.1076 | 0.9898 | 0.0027 | 0.9321 |
| 10 g/L | 3.5000 | 0.9866 | 0.072 | 0.9934 | 0.0017 | 0.9447 |
| 20 g/L | 1.8968 | 0.9743 | 0.0495 | 0.9652 | 0.0018 | 0.7635 |
| 30 g/L | 1.4226 | 0.9758 | 0.0244 | 0.9914 | 0.0004 | 0.9778 |
| 40 g/L | 1.119 | 0.9986 | 0.0158 | 0.9885 | 0.0002 | 0.9630 |

## References

Ahmad, M.A., Herawan, S.G., Yusof, A.A., 2014. Equilibrium, kinetics, and thermodynamics of Remazol Brilliant Blue R dye adsorption onto activated carbon prepared from pinang frond. ISRN Mechanical Engineering. Article ID 184265.

Annuar, M.S.M., Adnan, S., Vikineswary S., Chisti Y., 2009. Kinetics and energetics of azo dye decolorization by *Pycnoporus sanguineus*. Water, Air, Soil Pollu. 202, 179–188.

Antelo, F.S., Costa, J.A.V., Kalil, S.J., 2008. Thermal degradation kinetics of phycocyanin from *Spirulina platensis*. Biochem. Eng. J. 41, 43–47.

Bas, D., Dudak, F.C., Boyaci, I.H., 2007. Modeling and optimization IV: investigation of reaction kinetics and kinetic constants using a program in which artificial neural network (ANN) was integrated. J. Food Eng. 79: 1152–1158.

Brown, J.P., 1981. Reduction of polymeric azo and nitro dyes by intestinal bacteria. Appl. Environ. Microbiol. 41, 1283–1286.

Cao, J., Sanganyado, E., Liua, W., Zhang, W., Liu, Y., 2019. Decolorization and detoxification of Direct Blue 2B by indigenous bacterial consortium. J. Environ. Manage. 242, 229–237.

Carliell, C.M., Barclay, S.J., Naidoo, N., Buckley, C.A., Mulholland, D.A., Senior, E., 1995. Microbial decolourisation of a reactive azo dye under anaerobic conditions. Water SA. 21(1), 61–69.

Chang, J.S., Chou, C., Lin, Y.C., Lin, P.J., Ho, J.Y., Hu, T.L., 2001. Kinetic characteristics of bacterial azo-dye decolourization by *Pseudomonas luteola*. Water Res. 35(12), 2841–2850.

Dafale, N., Wate, S., Meshram, S., Nandy, T., 2008. Kinetic study approach of remazol black-B use for the development of two-stage anoxic–oxic reactor for decolorization/biodegradation of azo dyes by activated bacterial consortium. J. Hazard. Mater.159, 319–328.

Dubin, P., Wright, K.L., 1975. Reduction of azo dyes in cultures of *Proteus vulgaris*. Xenobiotics. 5, 563–571.

Eskandari, F., Shahnavaz, B., Mashreghi, M., 2019. Optimization of complete RB-5 azo dye decolorization using novel cold adapted and mesophilic bacterial consortia. J. Environ. Manage. 241, 91–98.

Houng, J.Y., Liau, J.S., 2006. Mathematical modeling of asymmetric reduction of ethyl 4-chloro acetoacetate by bakers’ yeast. Enzyme Microb. Technol. 38, 879–886.

Jain, K., Shah, V., Chapla, D., Madamwar, D., 2012. Decolorization and degradation of azo dye – Reactive Violet 5R by an acclimatized indigenous bacterial mixed cultures-SB4 isolated from anthropogenic dye contaminated soil. J. Hazard. Mater. 213– 214, 378– 386.

Lourenco, N.D., Novais, J.M., Pinheiro, H.M., 2006. Kinetic studies of reactive azo dye decolorization in anaerobic/aerobic sequencing batch reactors. Biotechnol. Lett. 28, 733–739.

Sponza, D.T., Isik, M., 2004. Decolorization and inhibition kinetic of direct black 38 azo dye with granulated anaerobic sludge. Enzyme Microb. Technol. 34, 147–158.

van der Zee F.P., Lettinga, G., Field, J.A., 2001. Azo dye decolorization by anaerobic granular sludge. Chemosphere. 44, 1169–1176.

Weber, E.J., Wolfe, L.N., 1987. Kinetic studies of reduction of aromatic azo compounds in anaerobic sediment water system. Environ. Toxicol. Chem. 6, 911–919.

Weber, E.J., 1991. Studies of benzidine based dyes in sediment-water system. Environ. Toxicol. Chem. 10, 609–618.

Wuhrmann, K., Mechnsner, K., Kappeler, T., 1980. Investigation on rate determining factors in microbial reduction of azo dyes. Eur. J. Appl. Microbiol. Biotechnol. 9, 325– 338.

Xie, X.H., Zheng, X.L., Yu, C.Z., Zhang, Q.Y., Wang, Y.Q., Cong, J.H., Liu, H., He, Z.J., Yang, B., Liu, J.S., 2020. HighDefficient biodegradation of refractory dye by a new bacterial flora DDMY1 under different conditions. Int. J. Environ. Sci. Technol. 17, 1491–1502

Yan, B., Du, C., Xu, M., Liao, W., 2012. Decolorization of azo dyes by a salt-tolerant *Staphylococcus cohnii* strain isolated from textile wastewater. Front. Environ. Sci. Eng. 6(6), 806–814.

